# Discovery of electrophilic degraders that exploit SNAr chemistry

**DOI:** 10.1101/2024.09.25.615094

**Authors:** Zhe Zhuang, Woong Sub Byun, Zuzanna Kozicka, Brendan G. Dwyer, Katherine A. Donovan, Zixuan Jiang, Hannah M. Jones, Dinah M. Abeja, Meredith N. Nix, Jianing Zhong, Mikołaj Słabicki, Eric S. Fischer, Benjamin L. Ebert, Nathanael S. Gray

## Abstract

Targeted covalent inhibition (TCI) and targeted protein degradation (TPD) have proven effective in pharmacologically addressing formerly ‘undruggable’ targets. Integration of both methodologies has resulted in the development of electrophilic degraders where recruitment of a suitable E3 ubiquitin ligase is achieved through formation of a covalent bond with a cysteine nucleophile. Expanding the scope of electrophilic degraders requires the development of electrophiles with tempered reactivity that enable selective ligase recruitment and reduce cross-reactivity with other cellular nucleophiles. In this study, we report the use of chemical moieties that enable nucleophilic aromatic substitution (S_N_Ar) reactions in the rational design of electrophilic protein degraders. Appending an S_N_Ar covalent warhead to several preexisting small molecule inhibitors transformed them into degraders, obviating the need for a defined E3 ligase recruiter. The S_N_Ar covalent warhead is versatile; it can recruit various E3 ligases, including DDB1 and CUL4 associated factor 11 (DCAF11), DDB1 and CUL4 associated factor 16 (DCAF16), and possibly others. The incorporation of an S_N_Ar covalent warhead into the BRD4 inhibitor led to the discovery of degraders with low picomolar degradation potency. Furthermore, we demonstrate the broad applicability of this approach through rational functional switching from kinase inhibitors into potent degraders.

## Main

Targeted protein degradation is a promising therapeutic strategy for eliminating disease-relevant proteins that are not addressable through conventional occupancy-driven pharmacology^1,2^. Protein degraders induce proximity between E3 ubiquitin ligases and target proteins, ultimately leading to ubiquitination and subsequent target depletion. A major class of protein degraders are proteolysis-targeting chimeras (PROTACs), and are bivalent molecules composed of ligands for the targeted protein and an E3 ubiquitin ligase, connected by a suitable linker^3,4^. However, the usually high molecular weight poses significant challenges for obtaining suitable pharmacokinetic (PK) properties, leading to an often challenging and lengthy drug development process^5^. In contrast, monovalent molecular glue degraders exhibit better drug-like properties as exemplified by the clinical success of thalidomide derivative oral drugs^6^. Nevertheless, discovery of monovalent molecular glues remains serendipitous, and optimizing their medicinal chemistry can be challenging^7^.

Both targeted covalent inhibition (TCI) and targeted protein degradation (TPD) have demonstrated their ability to drug targets that were previously considered to be ‘undruggable’^8^. Recently, these two approaches have been merged to enable the creation of small molecule degraders bearing an electrophilic group that can recruit a ligase through covalent bond formation^9^. These compounds were developed using two distinct and complementary approaches. In the first, electrophilic fragments that covalently labelled E3 ubiquitin ligases identified by mass spectrometry-based proteomics were used for the development of covalent PROTACs, including DCAF1, DCAF11, DCAF16, RNF114, and FEM1B^10–14^. In the second approach, cell-based screening of degraders bearing electrophiles and subsequent mechanism of action studies allowed discovery of degraders that function through the recruitment of DCAF11, DCAF16, and RNF126^15–18^. Electrophilic degraders that covalently recruit the E3 ligase may be advantageous because they can (1) maintain catalytic degradation turnover, (2) recruit a diverse range of E3 ligases that lack binding pockets that can be adequately engaged by reversible ligands, and (3) enhance potency with prolonged duration of action. To date there has been limited exploration of covalent degraders and much remains to be further studied about how to develop them and what may be their advantages and disadvantages relative to reversible degraders^9^.

Nucleophilic aromatic substitution (S_N_Ar) is one of the oldest reactions in organic chemistry. Despite its historical significance as the second most frequently used reaction in medicinal chemistry^19^, its application in the development of targeted covalent inhibitors and degraders has yet to be thoroughly explored. Covalent warheads using a thio-S_N_Ar reaction feature tunable electrophilicity and adjustable distance. Representative examples include the development of irreversible, cysteine-targeted inhibitors of peroxisome proliferator-activated receptors (PPARs), fibroblast growth factor receptor 4 (FGFR4), mitogen-activated protein kinase-activated protein kinase 2 (MK2), hepatitis C virus (HCV) NS5B polymerase, and other targets^20–24^. Although there have been advancements in mapping ligandable cysteines by electron-deficient halogenated aryl fragments^25^, there is no precedent to our knowledge for applying S_N_Ar chemistry in TPD.

Herein, we harness S_N_Ar chemistry for the design of electrophilic protein degraders. Minor modifications to the parental inhibitors through the incorporation of an S_N_Ar covalent warhead led to protein degradation. The tunable S_N_Ar covalent warheads can recruit various E3 ubiquitin ligases, including DDB1 and CUL4 associated factor 11 (DCAF11), DDB1 and CUL4 associated factor 16 (DCAF16), and possibly others. Attaching an S_N_Ar warhead onto a BRD4 inhibitor led to the discovery of BRD4 degraders with low picomolar potency, amongst the most potent degraders described so far. We showcase the versatility of this approach by integrating the S_N_Ar electrophile into a range of ATP-competitive kinase inhibitors to generate degraders targeting AURKA, AURKB, PTK2B, ITK, LIMK2, CDK4, CDK6, CDK12, and WEE1. These electrophilic kinase degraders with the identical S_N_Ar warhead exhibit distinct mechanisms compared to electrophilic BRD4 degraders.

### Results and discussion: Introducing an S_N_Ar covalent warhead into a BRD4 inhibitor triggered the gain-of-function transition into a potent BRD4 degrader

The bromodomain and extra-terminal domain (BET) family of proteins are promising therapeutic targets for cancer and inflammatory diseases^26,27^. We began our investigation of developing S_N_Ar-enabled degraders by selecting BRD4 as the protein of interest (POI), given the existence of monovalent and bivalent electrophilic BRD4 degraders that covalently label E3 ubiquitin ligases based on JQ1, the widely used ligand of the BET family of bromodomain proteins including BRD4^26^.

We quantitatively measured endogenous BRD4 protein levels in Jurkat cells using a HiBiT-BRD4 assay^28^, which employs CRISPR/Cas9-mediated fusion of an 11-amino acid peptide-tag to BRD4. Consistent with their previously reported potencies, the JQ1-based BRD4 PROTAC dBET6^28^ and a monovalent BRD4 electrophilic degrader, MMH2, containing a vinyl sulfonamide covalent warhead^15^ exhibited DC_50_ values of 21 nM and 12 nM, respectively (Figure 1B). We prepared our series of JQ1 analogues by installing an S_N_Ar covalent warhead into a solvent-exposed region of the parental inhibitor (Figure S1). Among them, ZZ7-16-073, a JQ1 analogue with a 2-fluoro-5-nitrobenzoic amide warhead (Figure 1A), exhibited potent BRD4 degradation, with DC_50_ and D_max_ values of 15 pM and 97%, respectively (Figure 1B).

**Figure 1.**
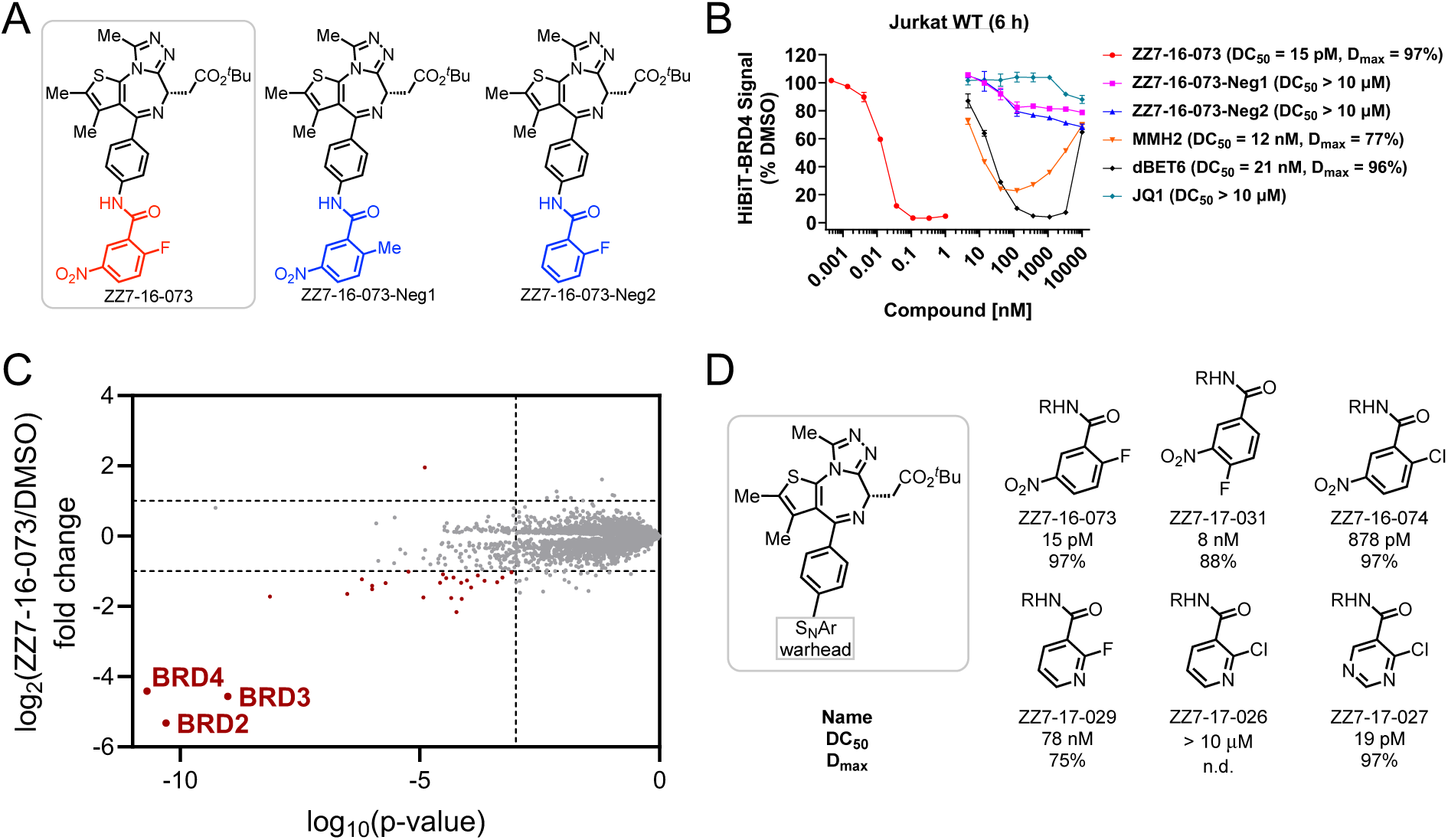
ZZ7-16-073 is a potent electrophilic BRD4 degrader dependent on the S_N_Ar covalent warhead. **A.** Structures of electrophilic BRD4 degrader ZZ7-16-073 and negative controls with slight chemical modifications that negate S_N_Ar reactivity. **B.** HiBiT-BRD4 results for Jurkat cells treated with the indicated compounds for 6 hours. **C.** Quantitative proteome-wide mass spectrometry in MOLT-4 cells after 3 hours treatment with 10 pM ZZ7-16-073. **D.** Structure-activity relationships (SAR) of electrophilic BRD4 degraders with an S_N_Ar covalent warhead.

To confirm the necessity of the S_N_Ar warhead, we synthesized two negative controls with slight chemical modifications that negate S_N_Ar reactivity. In ZZ7-16-073-Neg1, the fluoride leaving group is replaced with a methyl group (Figure 1A). ZZ7-16-073-Neg2 contains no nitro substituent (Figure 1A). Both negative controls lost the ability to degrade BRD4, suggesting the requirement of the S_N_Ar covalent warhead (Figure 1B and S2A). To examine the selectivity of ZZ7-16-073, we carried out quantitative proteome-wide mass spectrometry in both MOLT-4 and Jurkat cells after 3 hours treatment at a concentration of 10 pM to capture the most immediate protein level changes. We observed selective degradation of BRD2, BRD3, and BRD4, matching what has been reported for JQ1-based PROTACs (Figure 1C and S2D)^28^. We also found that ZZ7-16-073 exhibited low picomolar growth inhibition across diverse cancer cell lines (Figure S3).

We synthesized several analogues of ZZ7-16-073 to study structure-activity relationships (SAR) (Figure 1D). Moving the fluoride leaving group from the *ortho* to *para* position resulted in decreased DC_50_ values of 8 nM, suggesting that the geometry of the leaving group influenced the observed potency. The reactivity of the S_N_Ar warhead is crucial for potency, as replacing the fluoride leaving group with the less reactive chloride resulted in decreased potency (ZZ7-16-073 versus ZZ7-16-074; ZZ7-17-029 versus ZZ7-17-026). Given that the nitro group is considered a toxicophore in medicinal chemistry due to propensity to metabolism^29^, we explored several pyridine-containing S_N_Ar warheads, with ZZ7-17-029 exhibiting moderate potency (DC_50_ = 78 nM). Although its chloro analogue, ZZ7-17-026, failed to induce BRD4 degradation, increasing the reactivity of the warhead by introducing an electron-deficient moiety rescued BRD4 degradation; particularly noteworthy was the potency demonstrated by ZZ7-17-027 with the pyrimidine-containing S_N_Ar warhead, with a DC_50_ of 19 pM.

### The S_N_Ar warhead of ZZ7-16-073 covalently recruited DCAF16 Cys58

We next investigated the mechanism by which ZZ7-16-073 induces BRD4 degradation. Pretreatment with MLN-7243 (a ubiquitin-activating enzyme inhibitor), MLN-4924 (a NEDD8-activating enzyme inhibitor), or MG132 (a proteasome inhibitor) rescued BRD4 degradation in both the HiBiT-BRD4 degradation assay and Western blotting in Jurkat cells (Figure 2A and S4A). This indicates that ZZ7-16-073 promotes BRD4 degradation via the ubiquitin-proteasome system (UPS) ^30^. Co-treatment with the parental BRD4 binder JQ1 rescued BRD4 degradation, confirming the necessity of BET bromodomain binding (Figure 2A and S4A). ZZ7-16-073 induced degradation of first and second bromodomains of BRD4 (BRD4_BD1_ and BRD4_BD2_) with picomolar degradation potency (Figure S4B). In contrast, previous published BRD4 molecular glue degraders all preferentially degrade BRD4_BD2_^15,31^. We conducted a UPS-focused CRISPR degradation screen for both BRD4_BD1_ and BRD4_BD2_ to identify ubiquitin-related molecular machinery required for the degradation^32^. K-562 cells engineered to express Cas9 along with either the BRD4_BD1_ or BRD4_BD2_ reporter were transduced to express a single-guide RNA (sgRNA) library targeting ubiquitin-related genes^32^. The CRISPR knockout screening indicated that the knockout of DCAF16 and its associated components abrogated ZZ7-16-073’s BRD4-degrading potency in both isolated bromodomain reporters (Figure 2B). We validated the screening result via DCAF16 knockout HiBiT-BRD4 degradation assay (Figure 2C).

**Figure 2.**
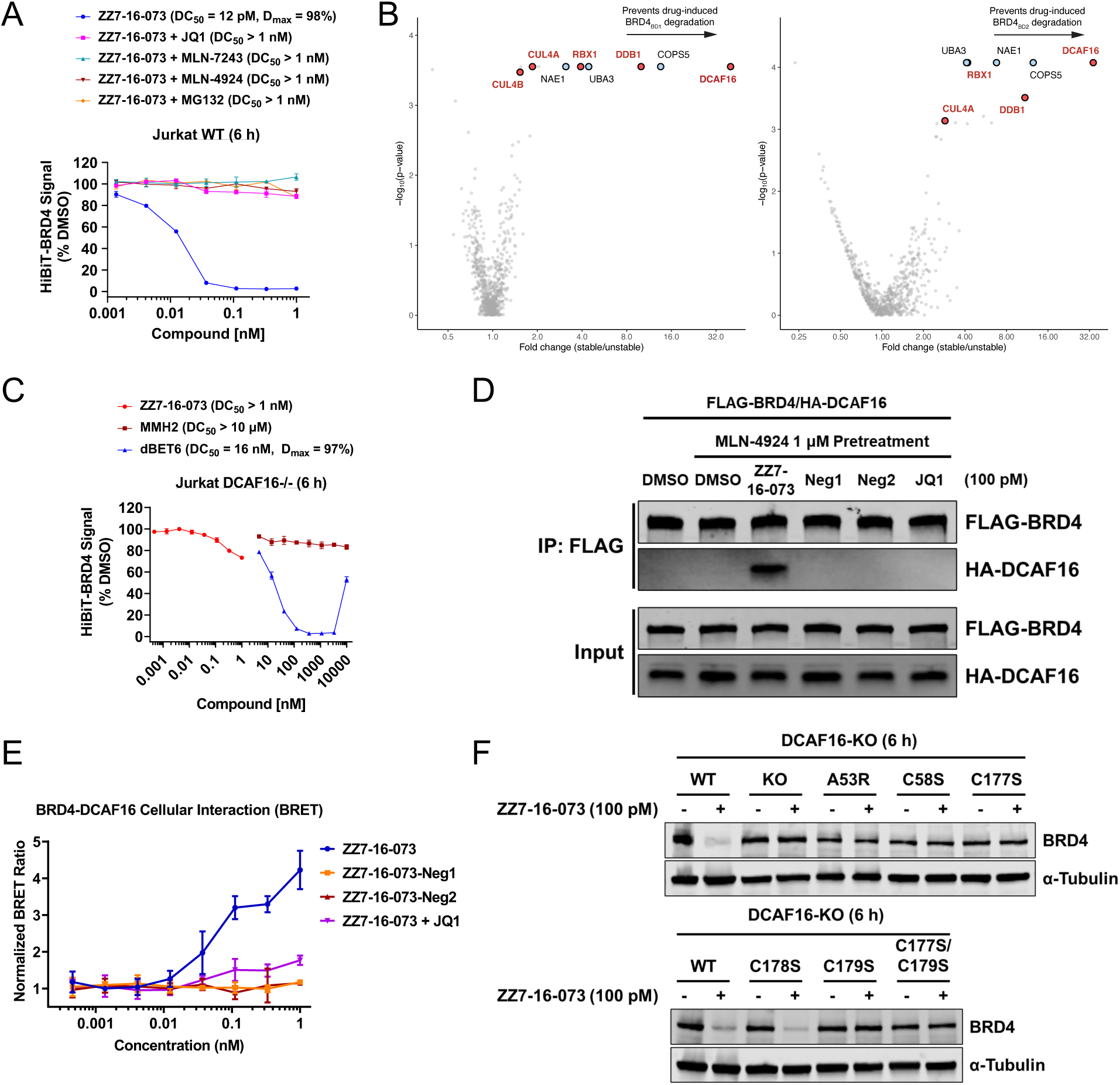
ZZ7-16-073-induced BRD4 degradation is dependent on DCAF16. **A.** HiBiT-BRD4 results for Jurkat cells treated with the indicated inhibitors for 1 hour and then ZZ7-16-073 for 6 hours. **B.** Ubiquitin-proteasome system (UPS)-focused CRISPR degradation screen for BRD4_BD1_-eGFP and BRD4_BD2_-eGFP stability in K562-Cas9 cells treated with 500 nM ZZ7-16-073 for 16 hours. **C.** HiBiT-BRD4 results for DCAF16-KO Jurkat cells treated with the indicated compounds for 6 hours. **D.** Co-immunoprecipitation of FLAG-tagged BRD4 and HA-tagged DCAF16 in the presence of the indicated compounds. **E.** NanoBRET assay suggesting the induced interaction between HaloTag-DCAF16 and Nanoluciferase-BRD4 fusion proteins with the indicated compounds. **F.** Western blots of BRD4 degradation in WT or DCAF16-KO and mutants K562 cells treated with 100 pM ZZ7-16-073 for 6 hours.

To confirm target engagement, we expressed FLAG-tagged BRD4 and HA-tagged DCAF16 in HEK293T cells. Immunoprecipitation of FLAG-tagged BRD4 co-enriched HA-tagged DCAF16 only in the presence of ZZ7-16-073 at 100 pM and not with the negative control compounds or JQ1, indicating that ZZ7-16-073 facilitates ternary complex formation between BRD4 and DCAF16 (Figure 2D). Co-immunoprecipitation with FLAG-tagged BRD4_BD1_ or BRD_BD2_ further confirmed the compound-dependent interaction with DCAF16 (Figure S4C). We also implemented a cellular NanoBRET assay to assess the induced interaction between HaloTag-DCAF16 and Nanoluciferase-BRD4 fusion proteins^33^. Only ZZ7-16-073 promoted proximity between DCAF16 and BRD4 in a dose-dependent manner, and this interaction was competitively inhibited by pretreatment with JQ1 (Figure 2E). We further confirmed the degradation was DCAF16-dependent by Western blotting in DCAF16 knockout K562 cells (Figure 2F). To identify the cysteine covalently labeled by ZZ7-16-073, we conducted a screening of BRD4 degradation using various DCAF16 mutant cell lines^15^. As a result, we identified Cys58 as exclusively necessary for ZZ7-16-073-induced BRD4 degradation (Figure 2F). Moreover, previous work showed Ala53, Cys177, and Cys179 were crucial for the E3 ligase function of DCAF16, in line with our observation (Figure 2F)^15^.

### Changing the attachment site of the S_N_Ar warhead altered the selectivity towards E3 ubiquitin ligase

To further define the degradation SAR for S_N_Ar-enabled BRD4 degraders, we identified ZZ7-16-078, a JQ1 analogue featuring the same S_N_Ar covalent warhead as ZZ7-16-073 but a different exit vector (Figure 3A). This compound exhibited moderate BRD4 degradation within 6 hours (DC_50_ = 459 nM and D_max_ = 58%) (Figure 3B). Shifting the fluoride leaving group from *ortho* to *para* position yielded ZZ7-17-033, which improved DC_50_ and D_max_ values of 25 nM and 87%, respectively (Figure 3B). This illustrated the importance of orientation for productive degradation using the S_N_Ar warhead. Both negative chemical controls with non-reactive moieties successfully rescued BRD4 degradation (Figure 3B and S5B). Pretreatment with MLN-7243, MLN-4924, or MG132 also abolished BRD4 degradation (Figure 3C). Quantitative proteome-wide mass spectrometry analysis in MOLT-4 and Jurkat cells showed that BRD2, BRD3, and BRD4 were the primary targets of degradation (Figure 3D and S5C).

**Figure 3.**
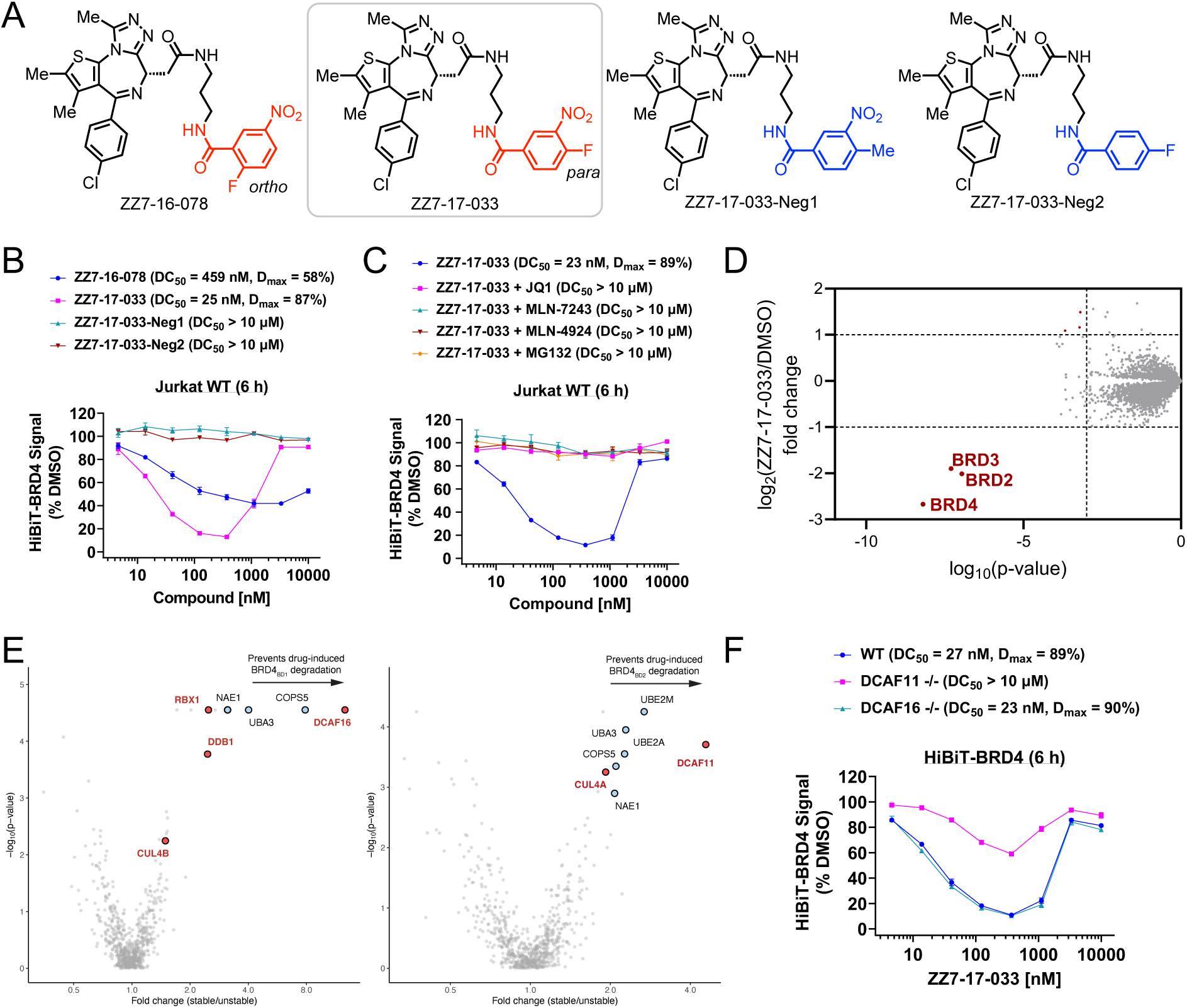
ZZ7-17-033-induced BRD4 degradation is primarily dependent on DCAF11. **A.** Structures of electrophilic BRD4 degrader ZZ7-16-078, ZZ7-17-033, and negative controls of ZZ7-17-033 with slight chemical modifications that negate S_N_Ar reactivity. **B.** HiBiT results for Jurkat cells treated with the indicated compounds for 6 hours. **C.** HiBiT-BRD4 results for Jurkat cells treated with the indicated inhibitors for 1 hours and then ZZ7-17-033 for 6 hours. **D.** Quantitative proteome-wide mass spectrometry in MOLT-4 cells after 3 hours treatment with 100 nM ZZ7-17-033. **E.** Ubiquitin-proteasome system (UPS)-focused CRISPR degradation screen for BRD4_BD1_-eGFP or BRD4_BD2_-eGFP stability in K562-Cas9 cells treated with 1 μM ZZ7-17-033 for 16 hours. **F.** HiBiT-BRD4 results for WT, DCAF11-KO, or DCAF1-KO Jurkat cells treated with ZZ7-17-033 for 6 hours.

To investigate the possibility that subtle alterations to the electrophile might have altered the BRD4 degradation mechanism, we performed a UPS-focused CRISPR degradation screen with ZZ7-17-033 to identify the E3 ligase responsible for BRD4 degradation. Surprisingly, while BRD4_BD1_ degradation required DCAF16, BRD4_BD2_ degradation required DCAF11 (Figure 3E). The BRD4 degradation by ZZ7-17-033 was nearly fully rescued in DCAF11 knockout Jurkat cells, indicating that DCAF11 is the primary E3 ligase driving the degradation (Figure 3F and S5D). Co-immunoprecipitation with FLAG-tagged BRD4 and a cellular NanoBRET assay as described above indicated that ZZ7-17-033 facilitated the formation of a ternary complex between BRD4 and DCAF11 (Figure S5E and S5F). Collectively, these findings demonstrate that attaching an identical covalent warhead through different exit vectors to the same parental inhibitor can switch the recruited E3 ubiquitin ligase^16,31^.

### Application of S_N_Ar chemistry to kinase degradation

To investigate the broader applicability of the S_N_Ar covalent warhead in designing electrophilic degraders for other POIs, we focused on kinases, for which there is an abundance of diverse inhibitors with a high potential for clinical impact. We paired an S_N_Ar warhead onto the multitargeted kinase inhibitor TL13-87 with different linker lengths, thereby creating a small library of potential promiscuous kinase degraders (Figure 4A). From this library, ZZ7-18-043 and ZZ7-18-045 both induced CDK12, AURKA, AURKB, PTK2B, ITK, and LIMK2 degradation in MOLT-4 cells treated for 3 hours at a concentration of 1 μM (Figure 4B). These kinases are among the primary targets of CRBN or VHL-based promiscuous kinase PROTACs that contain the same parental scaffold^34,35^. We conducted quantitative whole-cell global degradation proteomics to exclude the possibility that the observed degradation was a result of secondary effects or non-specific cytotoxicity^36^. A 3-hour treatment with 1 μM ZZ7-18-045 in MOLT-4 cells revealed CDK12, AURKA, AURKB, PTK2B, ITK, and LIMK2 as the primary degraded targets (Figure 4C). ZZ7-18-045-Neg, containing a non-reactive moiety, rescued degradation, indicating that the S_N_Ar covalent warhead is required for activity (Figure 4B). Pretreatment with the proteasome inhibitor bortezomib or MLN-7243 restored the degradation of AURKA, AURKB, and LIMK2 (Figure 4D). Whole-cell global proteomics with ZZ7-18-043 under the same conditions resulted in the downregulation of a wide range of proteins, likely attributed to the non-specific cytotoxicity associated with the electrophile (Figure S6A).

**Figure 4.**
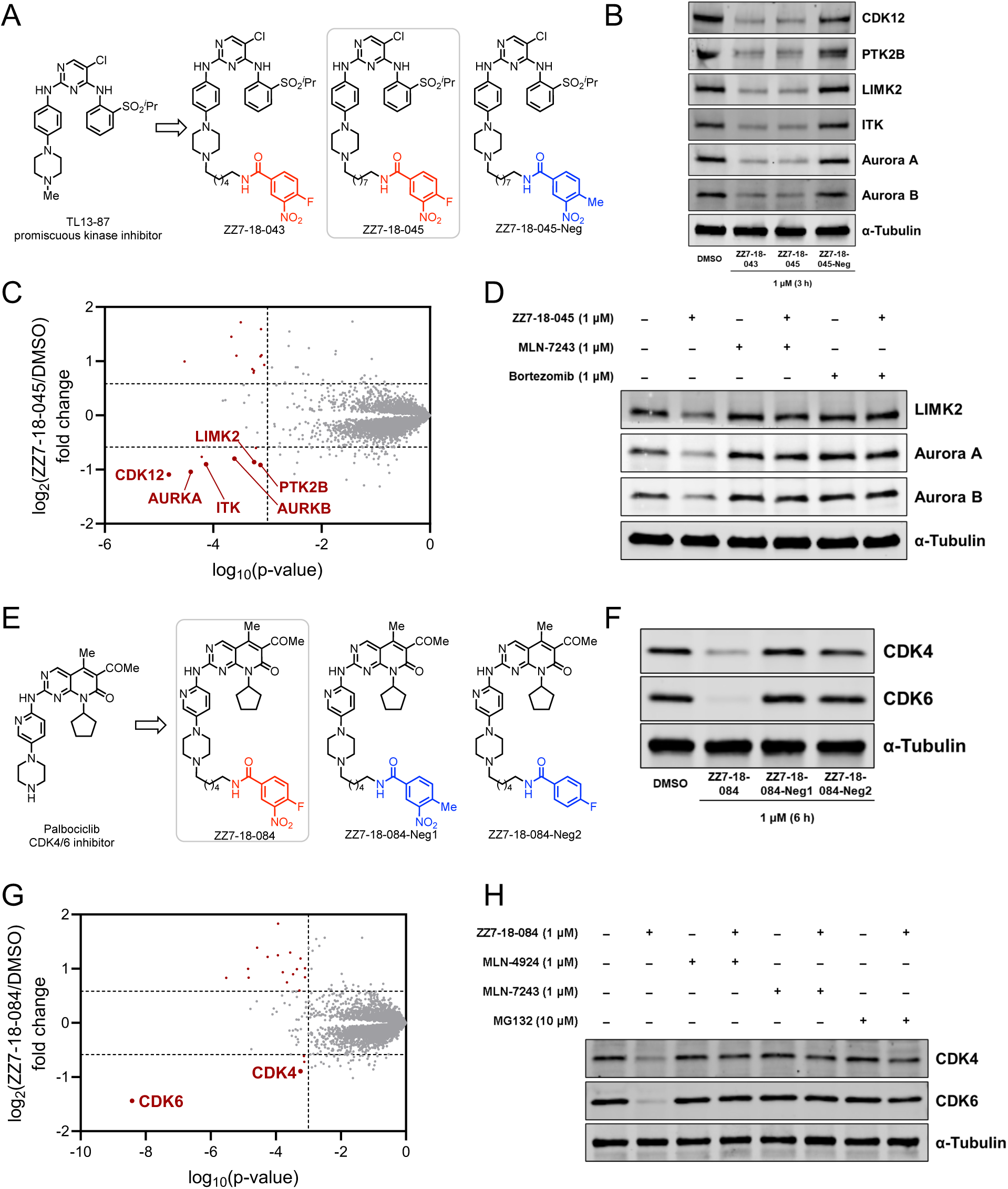
Development of electrophilic kinase degraders dependent on the S_N_Ar covalent warhead. **A.** Structures of electrophilic promiscuous kinase degraders ZZ7-18-043, ZZ7-18-045, and a negative control of ZZ7-18-045 with slight chemical modification that negate S_N_Ar reactivity. **B.** Western blots of kinases degradations in MOLT-4 cells treated for 3 hours with 1 μM indicated compounds. **C.** Quantitative proteome-wide mass spectrometry in MOLT-4 cells after 3 hours treatment with 1 μM ZZ7-18-045. **D.** Western blots of LIMK2, Aurora A, and Aurora B degradations in Jurkat cells treated with the indicated inhibitors for 1 hour and then ZZ7-18-045 for 3 hours. **E.** Structures of electrophilic CDK4/6 degrader ZZ7-18-084 and negative controls with slight chemical modifications that negate S_N_Ar reactivity. **F.** Western blots of CDK4 and CDK6 degradations in Jurkat cells treated with 1 μM indicated compounds for 6 hours. **G.** Quantitative proteome-wide mass spectrometry in MOLT-4 after 3 hours treatment with 1 μM ZZ7-18-084. **H.** Western blots of CDK4 and CDK6 degradations in Jurkat cells treated with the indicated inhibitors for 1 hour and then ZZ7-18-084 for 6 hours.

In addition to developing electrophilic degraders from a multitargeted kinase inhibitor, we have applied S_N_Ar chemistry to derive selective kinase degraders from kinase inhibitors. ZZ7-18-084 is derived from the FDA-approved CDK4/6 inhibitor palbociclib by attaching an S_N_Ar covalent warhead (Figure 4E)^37^. Whole-cell proteomics analysis in Jurkat cells conducted after a 3-hour treatment at a concentration of 1 μM showed that CDK4 and CDK6 were the primary degraded targets (Figure 4G and S6B). Moreover, ZZ7-18-084-induced CDK4 and CDK6 degradations are covalency- and UPS-dependent (Figure 4F and 4H). Surprisingly, neither DCAF11 nor DCAF16 was required for CDK4 and CDK6 degradation (Figure S6C). We also developed S_N_Ar-enabled electrophilic WEE1 and CDK12 degraders whose activity is covalency- and UPS-dependent (Figure S7 and S8)^38,39^. Efforts to develop a selective AURKA electrophilic degrader from the AURKA inhibitor Alisertib have proven unsuccessful (Figure S9)^40^. Taken together, we have engineered selective electrophilic kinase degraders by attaching an S_N_Ar covalent warhead to the parental inhibitors. Preliminary experiments indicate the S_N_Ar-enabled kinase degraders exploit distinct E3 ligase pathways than those found to promote S_N_Ar-enabled BRD4 degradation.

## Conclusion

We harnessed S_N_Ar chemistry for the design of electrophilic degraders, integrating advantages from both targeted covalent inhibition (TCI) and targeted protein degradation (TPD) fields. By synthesizing a small collection of JQ1 electrophile modified compounds and using BRD4 degradation as a screening platform we rapidly identified potent degraders. Compounds capable of inducing BRD4 degradation can be subsequently modified to investigate the structural features required to enable them to be effective degraders. Mechanistic studies indicated that these compounds induce degradation by recruiting ligases such as DCAF11 and DCAF16.

The versatility of the S_N_Ar warhead display approach is demonstrated through its chemical diversification, ability to recruit various E3 ubiquitin ligases, with improved degradation potency. The observation that we can toggle from DCAF16 to DCAF11 dependent degradation through altering the trajectory of the reactive group, similar to what was previously observed with acrylamide-based compounds^16^, indicates that the complementarity of the BRD4 DCAF protein-protein interface is a key selectivity determinant for the compounds. Notably, the DCAF11-dependent BRD4 degrader is distinct from previously reported examples in that it is capable of degrading isolated BD1 and does not function through an intramolecular BD1-BD2 glue mechanism^31^. We demonstrate the versatility of this approach through the incorporation of the S_N_Ar electrophile onto a variety of selective ATP-site kinase heterocycles to generate degraders of kinases such as AURKA, AURKB, PTK2B, ITK, LIMK2, CDK4, CDK6, CDK12, and WEE1. These electrophilic kinase degraders, despite sharing the same S_N_Ar warhead, activate unique degradation mechanisms compared to electrophilic BRD4 degraders, highlighting potential variability in the recruited E3 ligase based on the target protein. Identifying the responsible degradation pathways is of major interest and the subject of ongoing studies.

Until recently, PROTAC development has necessitated identifying small molecules that recruit E3 ubiquitin ligases, then synthesizing libraries of bivalent degraders, and finally confirming the feasibility of utilizing these E3 ligase recruiters. However, substantial medicinal chemistry optimization is required to improve cell membrane permeability and PK properties for these non-drug-like compounds. In this and previous work^15,16^, we have developed an alternative strategy whereby appending a minimal covalent warhead onto preexisting small molecule inhibitors converts them into degraders, obviating the need for a bespoke E3 recruiting element. Several other labs have recently reported similar efforts along these lines^18,41^. These electrophilic degraders can maintain catalytic degradation turnover and recruit a wider array of E3 ligases compared to those used in bivalent degraders. Crucially for downstream drug development, these prototype small molecules are druglike and have relatively low molecular weights.

Electrophilic degraders may have non-specific cytotoxicity and potential off-target effects, as commonly observed in targeted covalent inhibition. Optimal electrophiles ideally demonstrate moderate reactivity to minimize off-target toxicity and maximize the therapeutic window. To capture the immediate protein level changes induced by degraders, conducting characterization requires shorter timeframes and lower concentrations to minimize secondary effects and non-specific cytotoxicity. Quantitative proteome-wide mass spectrometry analysis of electrophilic degraders can offer valuable insights into their global selectivity profile, thus optimizing the therapeutic window.

This approach to electrophilic modification of small molecule inhibitors results in a functional shift from the protein inhibition to degradation by recruiting multiple E3 ubiquitin ligases. We envision that electrophilic modification strategies can extend beyond degradation, such as stabilizing tumor suppressors by recruiting deubiquitinases and activating apoptotic signaling by recruiting transcriptional activators.

---

**Figure S1.**
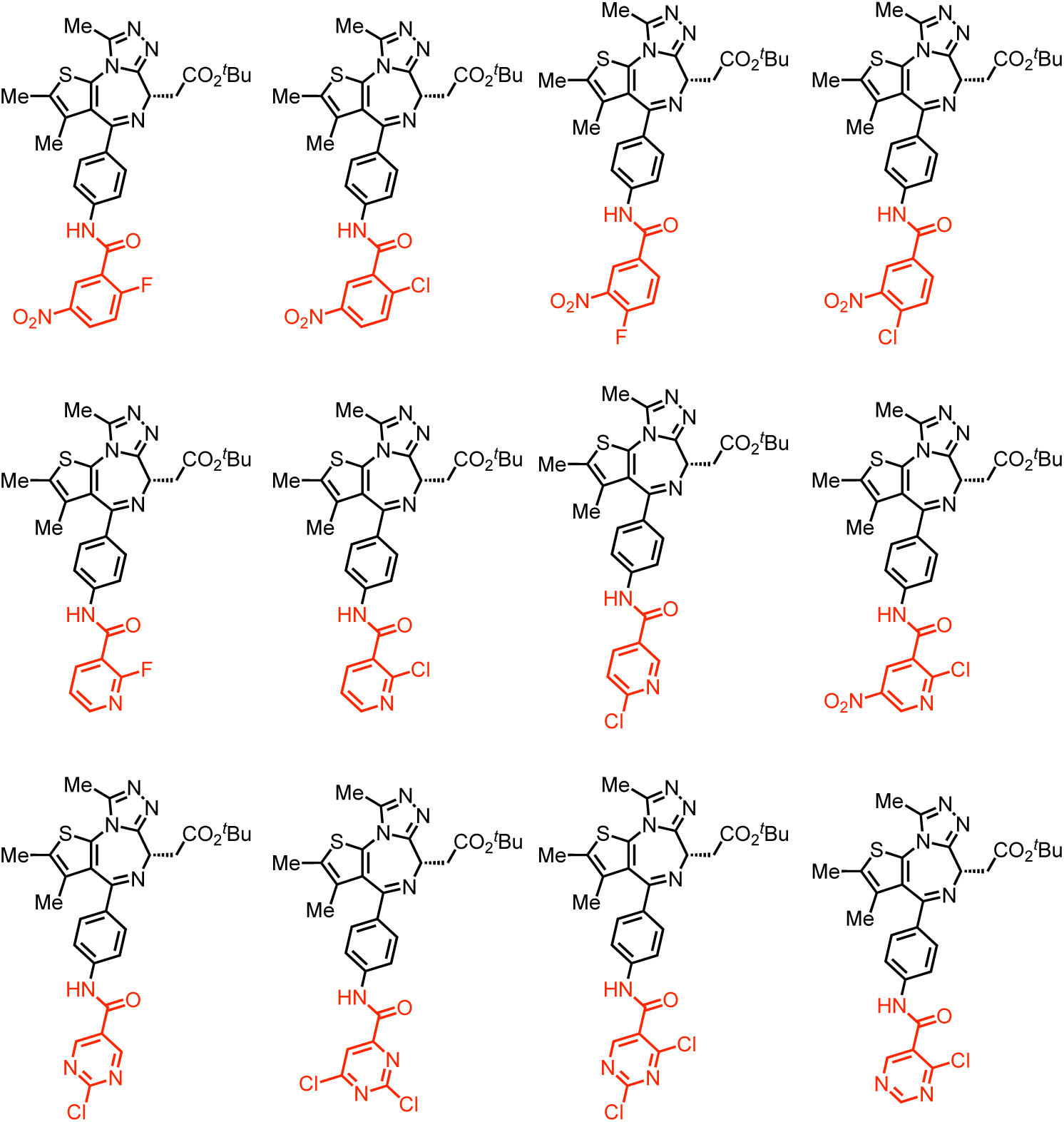
Structures of JQ1 analogues with an S_N_Ar covalent warhead.

**Figure S2.**
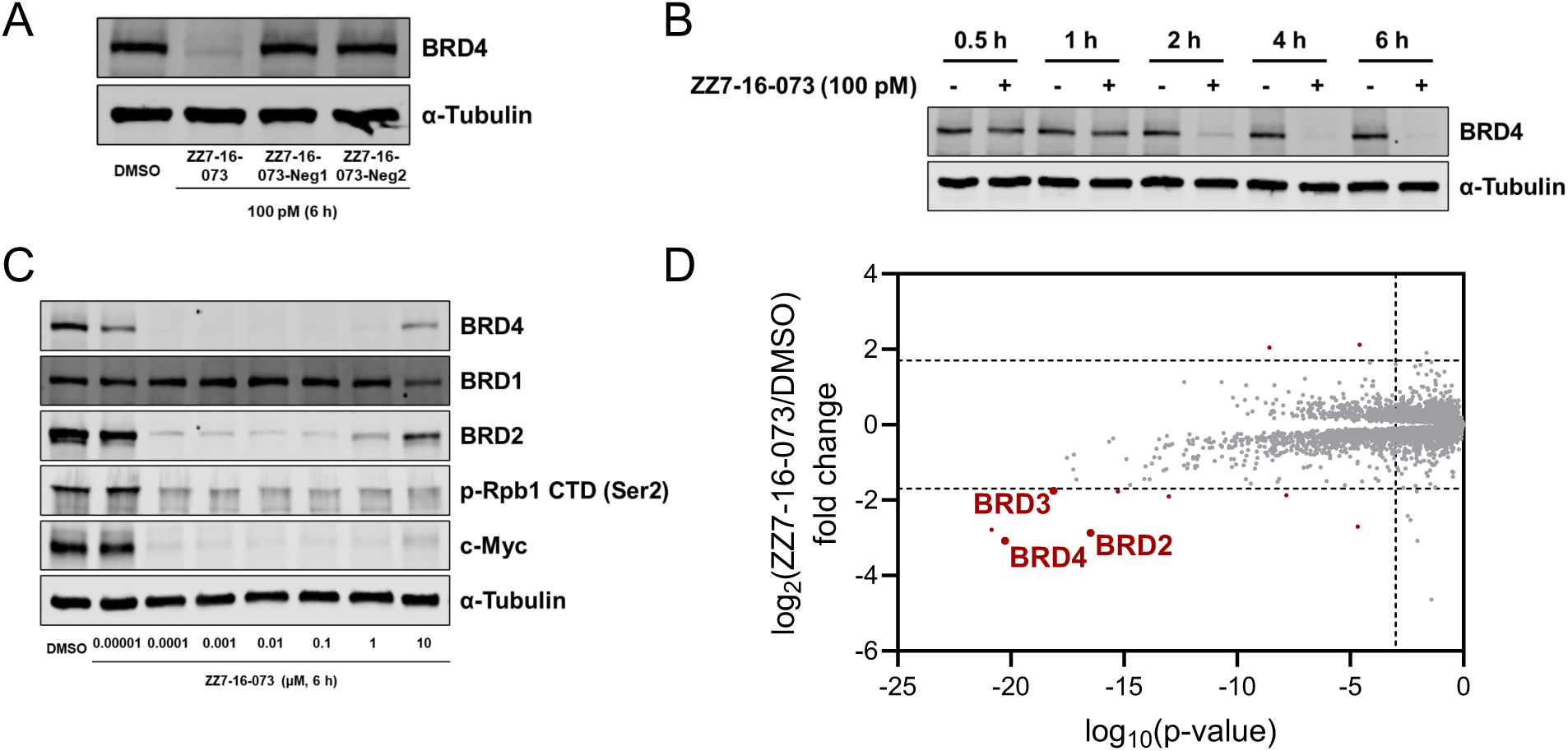
ZZ7-16-073 is a potent electrophilic BRD4 degrader dependent on the S_N_Ar covalent warhead. **A.** Western blots of BRD4 degradation in Jurkat cells treated with 100 pM indicated compounds for 6 hours. **B.** Western blots of BRD4 degradation in Jurkat cells treated with 100 pM ZZ7-16-073 for the indicated times. **C.** Western blots of BRD4, BRD2, and BRD1 degradations in Jurkat cells treated with the indicated concentrations of ZZ7-16-073 for 6 hours. **D.** Quantitative proteome-wide mass spectrometry in Jurkat cells after 3 hours treatment with 100 pM ZZ7-16-073.

**Figure S3.**
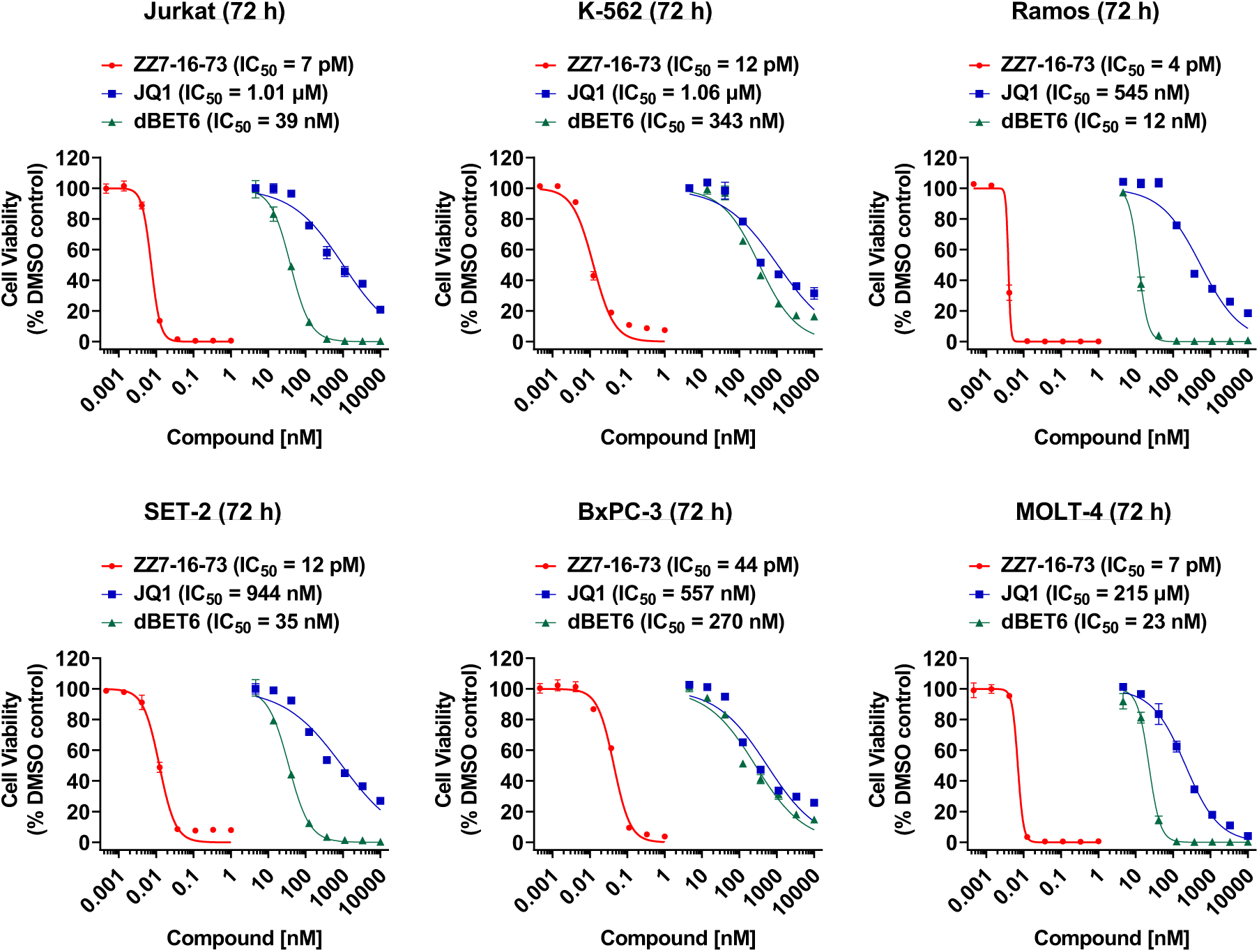
Growth inhibition by ZZ7-16-073, JQ1, and dBET6 across different cancer cell lines.

**Figure S4.**
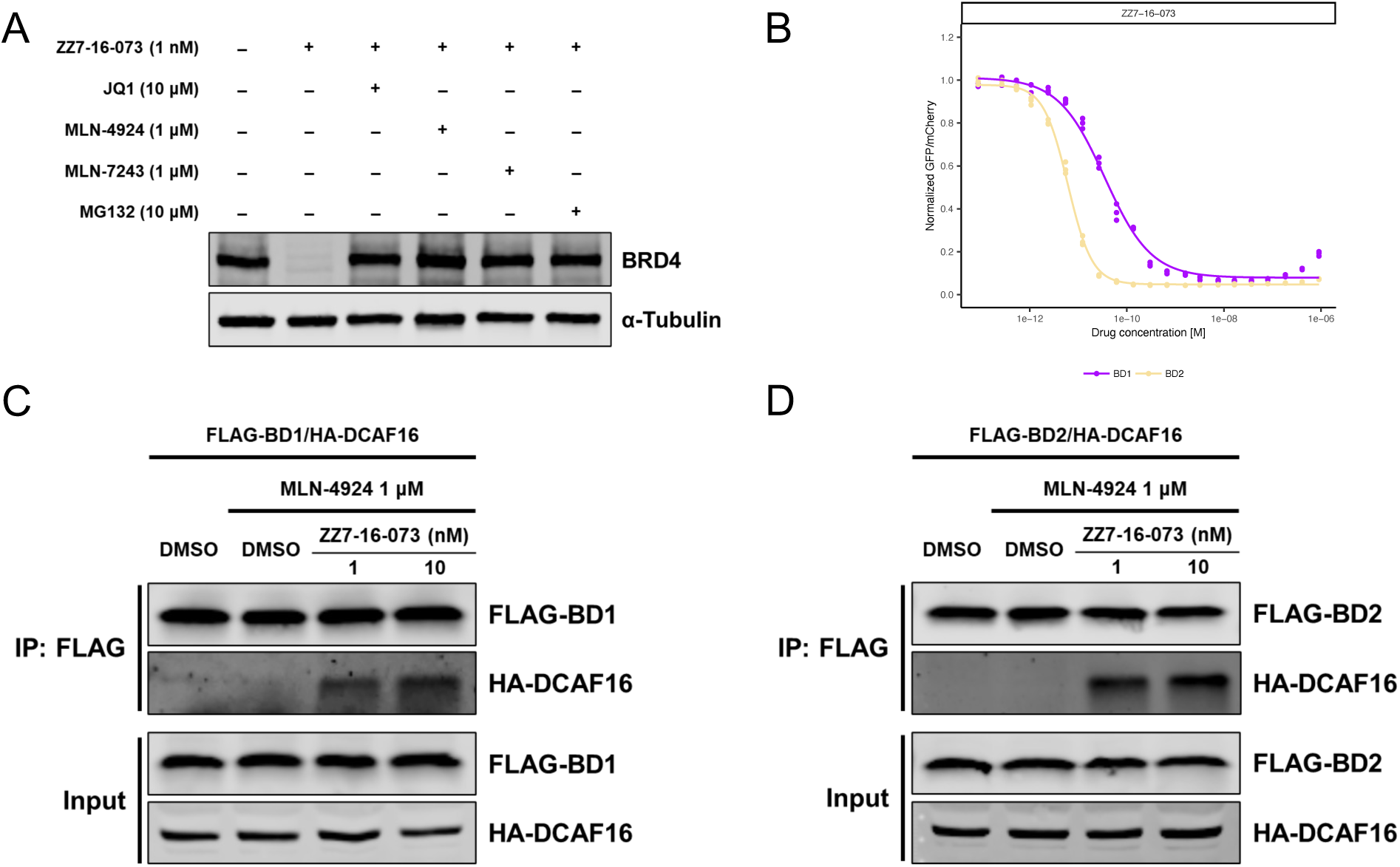
ZZ7-16-073-induced BRD4 degradation is dependent on DCAF16. **A.** Western blots of BRD4 degradation in Jurkat cells treated with the indicated inhibitors for 1 hour and then ZZ7-16-073 for 6 hours. **B.** Flow cytometry analysis of BRD4_BD1_-eGFP and BRD4_BD2_-eGFP degradation in K562 cells treated with ZZ7-16-073. **C.** Co-immunoprecipitation of FLAG-tagged BRD4_BD1_ and HA-tagged DCAF16 in the presence of ZZ7-16-073. **D.** Co-immunoprecipitation of FLAG-tagged BRD4_BD2_ and HA-tagged DCAF16 in the presence of ZZ7-16-073.

**Figure S5.**
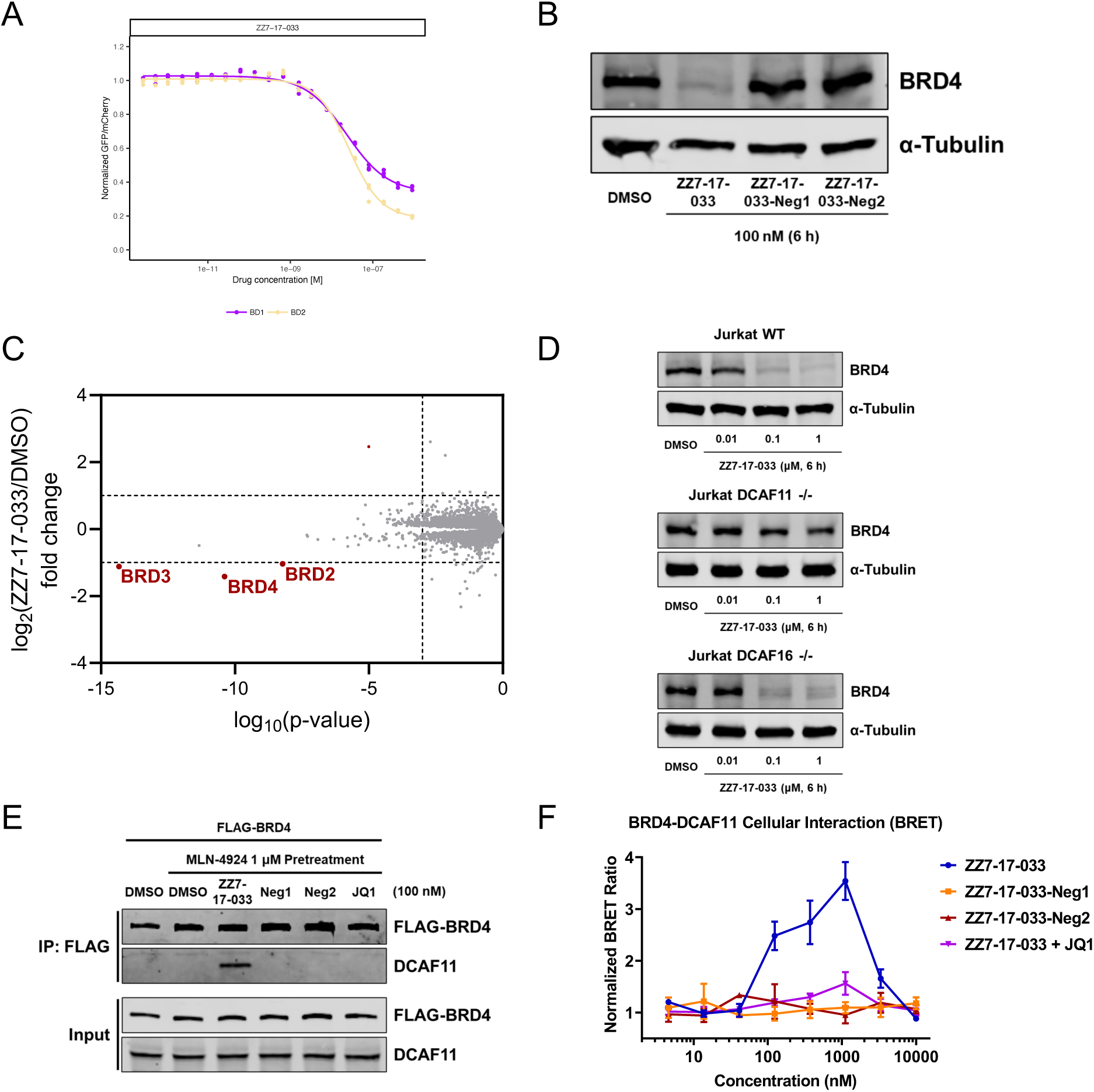
ZZ7-17-033-induced BRD4 degradation is primarily dependent on DCAF11. **A.** Flow cytometry analysis of BRD4_BD1_-eGFP and BRD4_BD2_-eGFP degradation in K562 cells treated with ZZ7-17-033. **B.** Western blots of BRD4 degradation in Jurkat cells treated for 6 hours with 100 nM indicated compounds. **C.** Quantitative proteome-wide mass spectrometry in Jurkat cells after 3 hours treatment with 100 nM ZZ7-17-033. **D.** Western blots of BRD4 degradation in WT, DCAF11-KO, or DCAF16-KO Jurkat cells treated with 100 nM ZZ7-17-033 for 6 hours. **E.** Co-immunoprecipitation of FLAG-tagged BRD4 and DCAF11 in the presence of the indicated compounds. **F.** NanoBRET assay suggesting the induced interaction between HaloTag-DCAF11 and Nanoluciferase-BRD4 fusion proteins with the indicated compounds.

**Figure S6.**
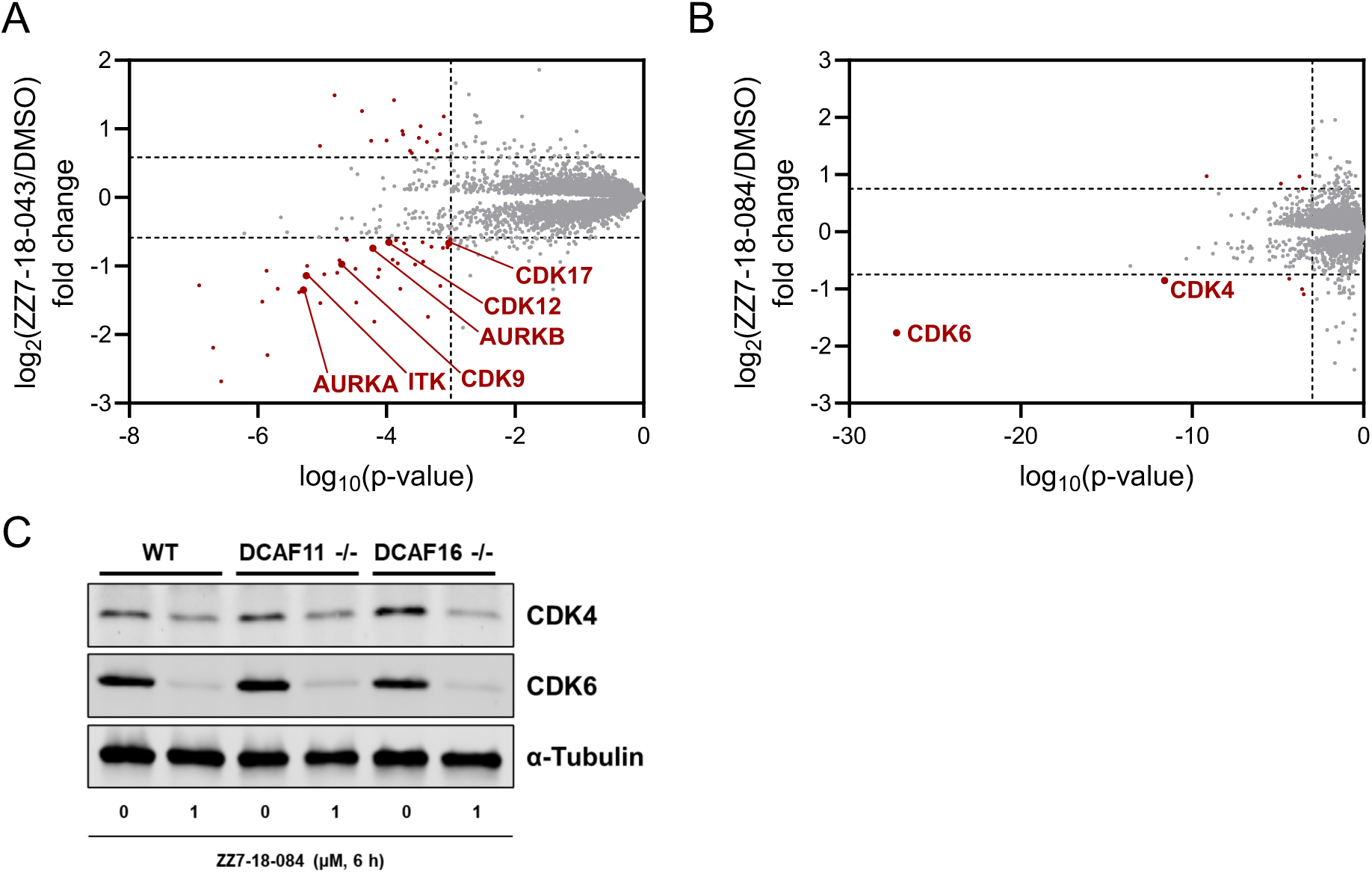
Development of electrophilic kinase degraders. **A.** Quantitative proteome-wide mass spectrometry in MOLT-4 cells after 3 hours treatment with 1 μM ZZ7-18-043. **B.** Quantitative proteome-wide mass spectrometry in MOLT-4 cells after 3 hours treatment with 1 μM ZZ7-18-084. **C.** Western blots of CDK4 and CDK6 degradations in WT, DCAF11-KO, or DCAF16-KO Jurkat cells treated for 6 hours with 1 μM ZZ7-18-084.

**Figure S7.**
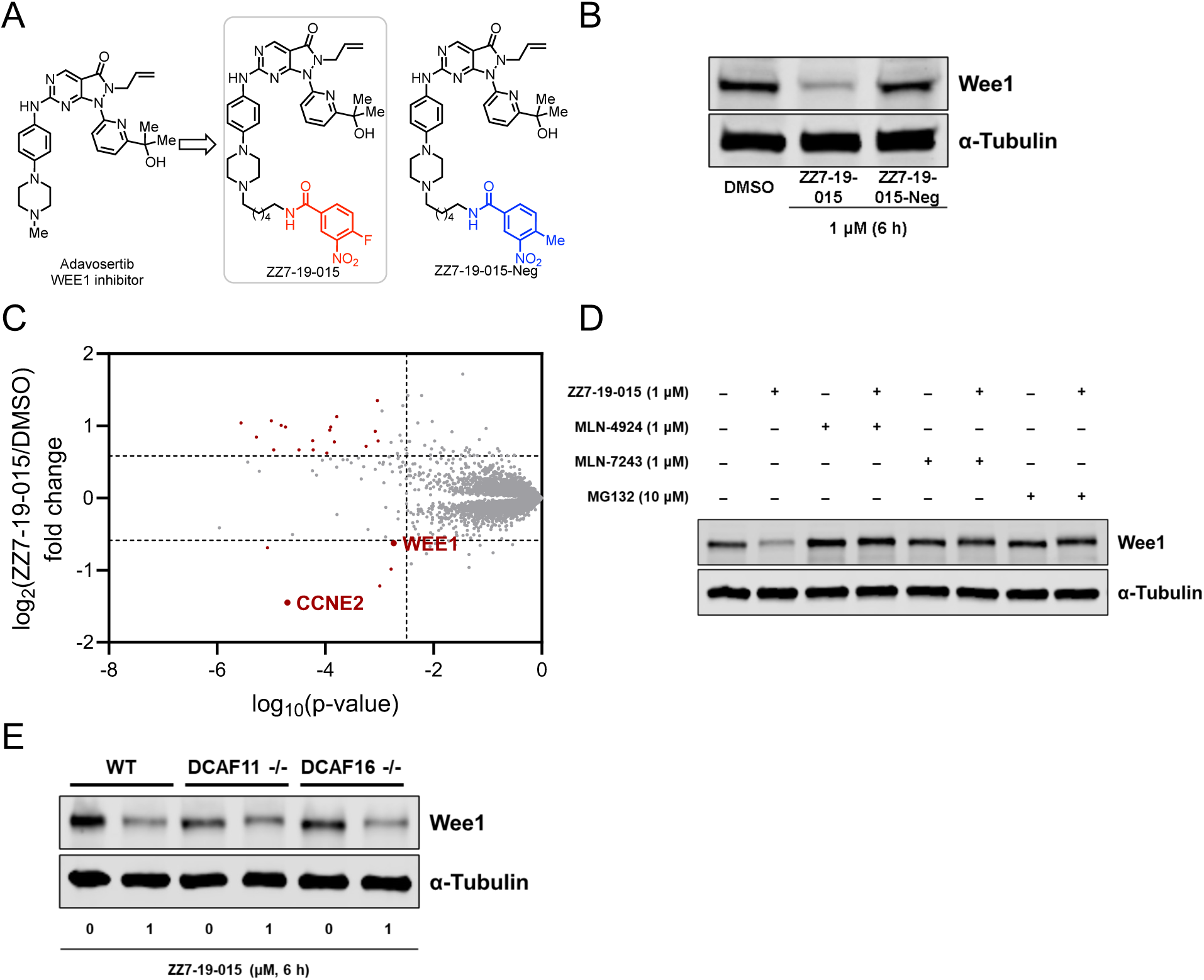
ZZ7-19-015 is an electrophilic WEE1 degrader dependent on the S_N_Ar covalent warhead. **A.** Structures of electrophilic WEE1 degrader ZZ7-19-015 and its negative control with slight chemical modification that negate S_N_Ar reactivity. **B.** Western blots of WEE1 degradation in Jurkat cells treated with 1 μM indicated compounds for 6 hours. **C.** Quantitative proteome-wide mass spectrometry in MOLT-4 cells after 3 hours treatment with 1 μM ZZ7-19-015. **D.** Western blots of WEE1 degradation in Jurkat cells treated with the indicated inhibitors for 1 hour and then ZZ7-19-015 for 6 hours. **E.** Western blots of WEE1 degradation in WT, DCAF11-KO, or DCAF16-KO Jurkat cells treated with 1 μM ZZ7-19-015 for 6 hours.

**Figure S8.**
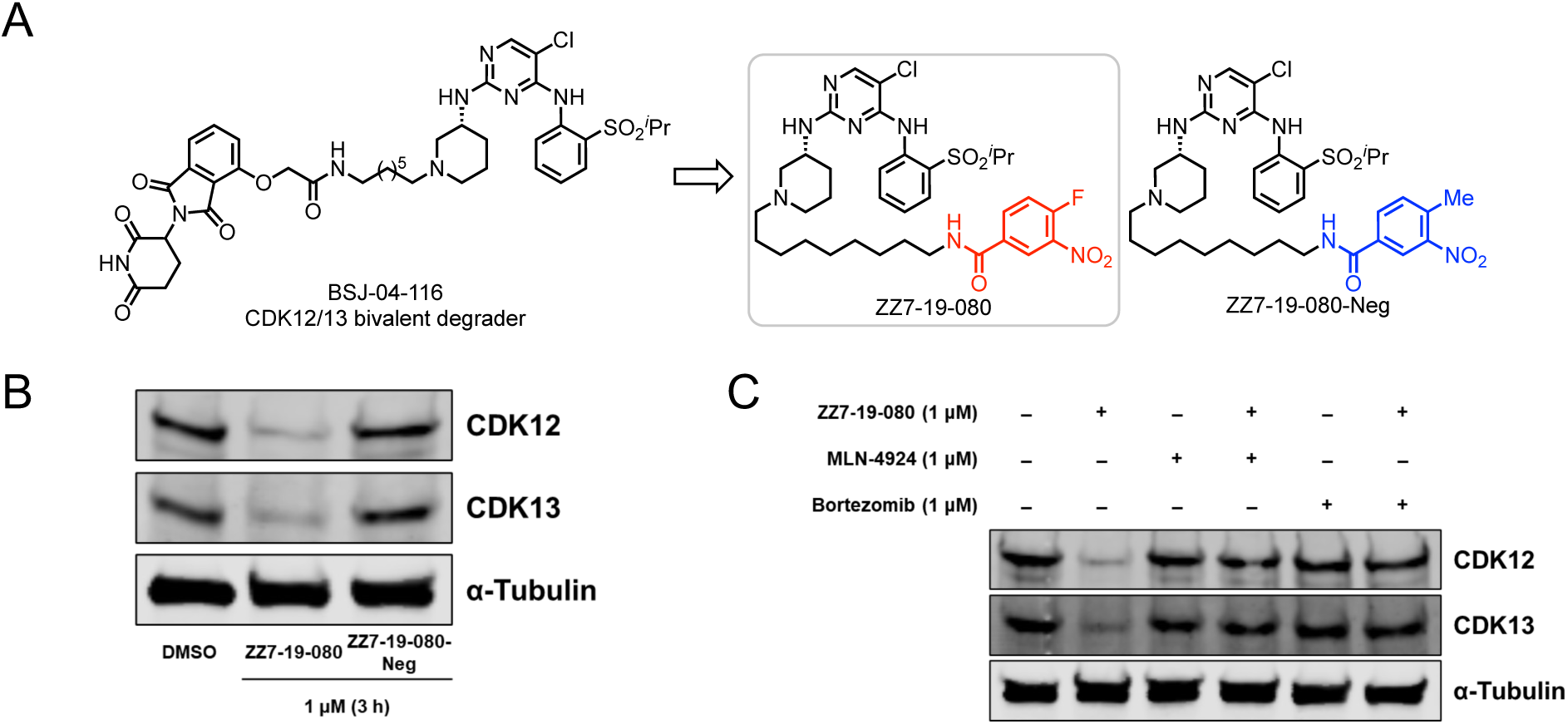
ZZ7-19-080 is an electrophilic CDK12/13 degrader dependent on the S_N_Ar covalent warhead. **A.** Structures of electrophilic CDK12/13 degrader ZZ7-19-080 and its negative control with slight chemical modification that negate S_N_Ar reactivity. **B.** Western blots of CDK12 and CDK13 degradations in Jurkat cells treated with 1 μM indicated compounds for 3 hours. **C.** Western blots of CDK12 and CDK13 degradation in Jurkat cells treated with the indicated inhibitors for 1 hour and then ZZ7-19-080 for 3 hours.

**Figure S9.**
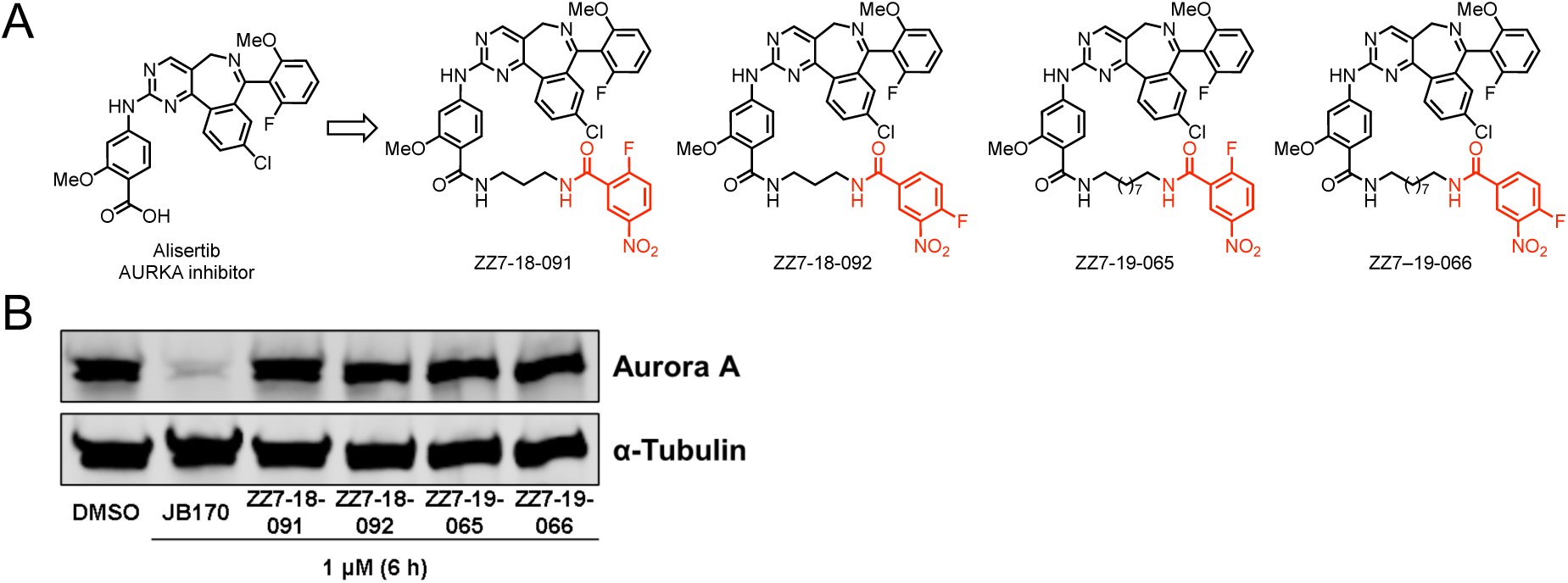
AURKA electrophilic modifiers with an S_N_Ar covalent warhead didn’t lead to AURKA degradation. **A.** Structures of AURKA electrophilic modifiers with an S_N_Ar covalent warhead. **B.** Western blots of AURKA degradation in Jurkat cells treated for 6 hours with 1 μM indicated compounds.

## Methods

### General information for biological assay

Roswell Park Memorial Institute (RPMI) 1640 medium and Dulbecco’s modified Eagle’s medium (DMEM), Fetal bovine serum (FBS), penicillin–streptomycin (10,000 units/mL sodium penicillin G and 10,000 μg/mL streptomycin), trypsin–EDTA solution (1×), and phosphate-buffered saline (PBS; 1×) were purchased from Gibco Invitrogen Corp. (Grand Island, NY, USA). JQ1, dBET6, MLN-7243, MLN-4924, MG132, Bortezomib were purchased from MedChemExpress (Monmouth Junction, NJ, USA). All other chemicals were purchased from Sigma–Aldrich (St. Louis, MO, USA), unless indicated otherwise.

### Mammalian cell culture

Human leukemia (Jurkat, K-562, Ramos, MOLT-4), pancreatic cancer (BxPC-3), and a kidney epithelial (HEK293T) cell lines were obtained from the American Type Culture Collection (ATCC, Manassas, VA, USA). A leukemia (SET-2) cell line was obtained from the Leibniz Institute DSMZ (Braunschweig, Germany). Cells were cultured in medium (RPMI 1640 for Jurkat, K-562, Ramos, MOLT-4, BxPC-3, SET-2 cells; DMEM medium for HEK293T cells) supplemented with 10% heat-inactivated FBS, 100 units/mL penicillin, 100 µg/mL streptomycin, and 0.25 µg/mL amphotericin B. Cells were incubated at 37 °C with 5% CO_2_ in a humidified atmosphere. Mycoplasma testing was performed monthly using the MycoAlert mycoplasma detection kit (Lonza, Basel, Switzerland) and all lines were negative. The human HEK293T cell lines were provided by the Genetic Perturbation Platform, Broad Institute and the K562-Cas9 by Z. Tothova (Dana-Farber Cancer Institute). HEK293T and K562-Cas9 cell lines were mycoplasma-negative and authenticated by STR profiling. HEK293T cells were cultured in Dulbecco’s modified Eagle’s medium (DMEM (Gibco) and K562-Cas9 in RPMI (Gibco), with 10% fetal bovine serum (FBS) (Invitrogen), glutamine (Invitrogen) and penicillin–streptomycin (Invitrogen) at 37 °C and 5% CO_2_.

### Generation of HiBiT-BRD4 cells

Introduction of a HiBiT coding sequence into the endogenous BRD4 locus in Jurkat cells was done via CRISPR-Cas9 genome editing. ALT-R CRISPR RNA (crRNA) and trans-activating CRISPR RNA (tracrRNA) (Integrated DNA Technologies, IDT) were resuspended in Nuclease-Free Duplex Buffer (IDT) at a concentration of 200 µM each. Equal volumes of crRNA and tracrRNA were mixed (final concentration of 100 µM each) and heated for 5 minutes at 95 °C. After heating, the complex was gradually cooled to room temperature. The oligo complex was then incubated at room temperature for 20 minutes with the Cas9 Nuclease V3 (IDT) to form the ribonucleoprotein (RNP) complex. The double-stranded DNA HDR template (HiBiT-BRD4 with extensions, see below), the RNP complex, and an electroporation enhancer (IDT) were then electroporated into Jurkat cells using an Amaxa 4D-Nucleofector (Lonza). Electroporated cells were transferred to medium with HRD enhancer (IDT). Single cells were subsequently isolated via fluorescence activated cell sorting, and HiBiT expression from individual clones was detected with the Nano-Glo HiBiT Lytic Detection System (Promega)^16^.

>crRNA sequence

ACTAGCATGTCTGCGGAGAG

>HDR donor sequence (+) CATTACTGGCAGATTTCTCAATCTCGTCCCAGGGCCGCTCTCCGCAGAGCCAGAACT

CCCGCTAATCTTCTTGAACAGCCGCCAGCCGCTCACCATGCTAGTGATCCCATCACATTCTTCACCAGGCACTCTA

>HDR donor sequence (-) TAGAGTGCCTGGTGAAGAATGTGATGGGATCACTAGCATGGTGAGCGGCTGGCGGCTGTTCAAGAAGATTAGCGGGAGTTCTGGCTCTGCGGAGAGCGGCCCTGGGACGAGATTGAGAAATCTGCCAGTAATG

### Generation of DCAF11 and DCAF16 knockout cells

CRISPR-Cas9 genome editing was used to knock out DCAF11 or DCAF16 in HiBiT-BRD4 Jurkat cells. crRNA and tracrRNA were resuspended in Nuclease-Free Duplex Buffer at mixed as described above. The oligo complex was subsequently incubated at room temperature with Cas9 Nuclease V3 (IDT) for 20 minutes to form the RNP complex. The RNP complex and electroporation enhancer (IDT) were electroporated into HiBiT-BRD4 Jurkat cells using an Amaxa-4D-Nucleofector (Lonza). Electroporated cells were transferred to medium with HDR enhancer (IDT). Single cells were isolated as above. Clones in which DCAF11 had been successfully knocked out were identified by immunoblotting for DCAF11^16^.

>crDNA sequence for DCAF11 knockout

TCGCGGAACAGCAGCAGTGC

>crDNA sequence for DCAF16 knockout

TCTGACAAGTGGTCAGGAGA

Genomic DNA from candidate knockout clones was purified (Machery Nagel) and used as template for PCR amplification using DNA oligos that anneal around the Cas9 cutting site. The resulting PCR products were gel-purified and submitted for next-generation sequencing (Quintara). Both knockout clones reported showed disruption of both alleles. The oligos used were as follows:

>DCAF16_fwd

TCCCTACACGACGCTCTTCCGATCTagtgttagtcagtgttctgctggcttag

>DCAF16_rev

GTTCAGACGTGTGCTCTTCCGATCTcaagactctcaagaggcgataagttggg

>DCAF11_fwd

TCCCTACACGACGCTCTTCCGATCTgctttcaccaaacccaaggaggtgacag

>DCAF11_rev

GTTCAGACGTGTGCTCTTCCGATCTgccagatccacatcttcatc

### HiBiT-BRD4 assay

Endogenous BRD4 protein levels were evaluated using the Nano-Glo HiBiT Lytic Detection System (Promega, Madison, WI, USA). Briefly, 2 x 10^4^ HiBiT-BRD4 Jurkat cells were seeded in 384-well plates and then cells were incubated with the indicated concentrations of compounds. After 6 h, the plates were subjected to Nano-Glo HiBiT Lytic Detection System Glo as described in manufacturer’s manual. The HiBiT-BRD4 assays were performed in biological triplicate. IC_50_ values were determined using a non-linear regression curve fit in GraphPad PRISM 9.5.1.

### Western blotting analysis

Total cells lysates were prepared in 2× sample loading buffer (i.e., 250 mM Tris-hydrochloride: pH 6.8, 4% sodium dodecyl sulfate, 10% glycerol, 0.006% bromophenol blue, 2% β-mercaptoethanol, 50 mM sodium fluoride, and 5 mM sodium orthovanadate). The samples with cell lysates were boiled for 5-8 min at 95 °C. The protein concentrations of the cell lysates were quantified using the BCA method and a BCA Protein Assay Kit (Thermo Fisher Scientific, Waltham, MA, USA). Equal amounts of protein were subjected to 4-20% sodium dodecyl sulfate-polyacrylamide gel electrophoresis and transferred to polyvinylidene fluoride membranes (Millipore, Bedford, MA, USA) activated with 100% methanol. The membranes were blocked using Intercept® (TBS) Blocking Buffer (LI-COR Biosciences, Lincoln, NE, USA), and subsequently probed with appropriate primary antibodies [anti-BRD4 (cat. no. ab128874; Abcam, Cambridge, UK), Anti-FLAG M2 (cat. no. F1804; Sigma-Aldrich), Anti-HA-Tag (cat. no. 3724; Cell Signaling Technology), anti-BRD1 (cat. no. sc-398226; Santa Cruz Biotechnology, Dallas, TX, USA), anti-BRD2 (cat. no. 5848; Cell Signaling Technology, Danvers, MA, USA), anti-phospho-Rpb1 CTD (Ser2) (cat. no. 04-1571; Sigma-Aldrich), anti-c-Myc (cat. no. 5605; Cell Signaling Technology), anti-α-Tubulin (cat. no. 3873; Cell Signaling Technology), anti-DCAF11 (cat. no. NBP2-92244; Novus Biologicals, Centennial, CO, USA), anti-CDK12 (cat. no. VMA00874; Bio-Rad Laboratories, Hercules, CA, USA), Anti-PTK2B (cat.no. 67141-1-Ig; Proteintech, Rosemont, IL, USA), anti-LIMK2 (cat. no. 3845; Cell Signaling Technology), anti-ITK (cat. no. 2380; Cell Signaling Technology), anti-Aurora A (cat. no. 91590; Cell Signaling Technology), anti-Aurora B (cat. no. 28711; Cell Signaling Technology), anti-CDK4 (cat. no. ab108355; Abcam), anti-CDK6 (cat. no. 3136; Cell Signaling Technology), anti-Wee1 (cat. no. 13084; Cell Signaling Technology), anti-CDK13 (cat. no. ABE1860; Sigma-Aldrich)] at 4 °C overnight and then incubated with IRDye 800-labeled goat anti-rabbit IgG (LI-COR Biosciences, cat. no. 926-32211) or IRDye 680RD goat anti-Mouse IgG (LI-COR Biosciences, cat. no. 926-68070) secondary antibodies at room temperature for 1 hour. After washing the membranes with PBS for 30 min, the membranes were detected on Li-COR Odyssey CLx system.

### Co-immunoprecipitation

HEK293T cells were seeded into 6-well plate (3 × 10^5^ cells/well), cultured overnight, and transfected with 1.5 μg FLAG-tagged and 1.5 μg HA-tagged plasmids using TransIT-LT1 transfection reagents (Mirus Bio, Madison, WI, USA). The transfected cells were cultured for another 36 h, pretreated with 1 μM MLN-4924, and co-treated with either compound or DMSO for 2 h before collection. The cells were collected and lysed in Pierce IP Lysis Buffer (Thermo Fisher Scientific) with cOmplete Mini Protease Inhibitor Cocktail (Roche, Basel, Switzerland) for 30 min on ice and centrifuged for 30 min at 4 °C to remove the insoluble fraction. For immunoprecipitation, 20 μL of pre-cleaned anti-FLAG M2 magnetic beads (Sigma-Aldrich) were added to the lysates. The beads–lysate mix was incubated at 4 °C for overnight on a rotator. Beads were magnetically removed and washed three times with PBS, and the FLAG-Tagged protein was competitively eluted using 3X FLAG Peptide (ApexBio Technology, Houston, TX, USA). Immunoblotting was carried out as previously described.

### NanoBRET assay for BRD4-DCAF interaction

HEK293T cells were seeded into 6-well plate (8 × 10^5^ cells/well), cultured overnight, and transfected with 2 μg HaloTag-DCAF and 0.1 μg NanoLuc-BRD4 plasmids using TransIT-LT1 transfection reagents (Mirus Bio). The cells were incubated overnight before replating in an opaque white 384-well plate at 2 x 10^5^ cells/mL in FluoroBrite DMEM (Gibco) with 10% FBS. Cells were treated with 0.1 μM HaloTag NanoBRET 618 ligand (Promega) or DMSO vehicle upon plating. The next day, the cells were pretreated for two hours with 1 μM MLN-4924 and then treated with the indicated compounds in triplicate for 2 hours. For JQ1 rescue experiments, cells were treated with 10 μM JQ1 for 1 hour after pretreatment with MLN-4924. A mixture containing 1:100 NanoGlo substrate:FluoroBrite DMEM was then added to each well and the plate was immediately read on a PHERAStar plate reader (BMG LABTECH, Cary, NC, USA) using a LUM 610-LP 450-80 optical module.

### Cell viability assay (CellTiter-Glo assay)

Cell viability was evaluated using the CellTiter-Glo assay (Promega, Madison, WI, USA). Briefly, cells were seeded in 384-well plates and then cells were incubated with the indicated concentrations of compounds. After 72 h, the plates were subjected to CellTiter-Glo as described in manufacturer’s manual. The proliferation assays were performed in biological triplicate. IC_50_ values were determined using a non-linear regression curve fit in GraphPad PRISM 9.5.1.

### Sample preparation for quantitative LFQ quantitative mass spectrometry

#### MOLT-4 proteomics

Cells were lysed by addition of lysis buffer (8 M Urea, 50 mM NaCl, 50 mM 4-(2-hydroxyethyl)-1-piperazineethanesulfonic acid (EPPS) pH 8.5, Protease and Phosphatase inhibitors) and homogenization by bead beating (BioSpec) for three repeats of 30 seconds at 2400 strokes/min. Bradford assay was used to determine the final protein concentration in the clarified cell lysate. Fifty micrograms of protein for each sample was reduced, alkylated and precipitated using methanol/chloroform as previously described^42^ and the resulting washed precipitated protein was allowed to air dry. Precipitated protein was resuspended in 4 M urea, 50 mM HEPES pH 7.4, followed by dilution to 1 M urea with the addition of 200 mM EPPS, pH 8. Proteins were digested with the addition of LysC (1:50; enzyme:protein) and trypsin (1:50; enzyme:protein) for 12 h at 37 °C. Sample digests were acidified with formic acid to a pH of 2-3 before desalting using C18 solid phase extraction plates (SOLA, Thermo Fisher Scientific). Desalted peptides were dried in a vacuum-centrifuged and reconstituted in 0.1% formic acid for liquid chromatography-mass spectrometry analysis.

#### Jurkat proteomics

Treated cell pellets were washed twice with TBS (50 mM Tris, pH 8.5, and 150 mM NaCl) and then thermally denatured in residual TBS for 5 min at 95 °C and then cooled to room temperature. Fresh Lysis Buffer (8 M urea, 150 mM NaCl, and 100 mM HEPES, pH 8.0, in MS-grade water) was then added and the lysates were homogenized using needle ultrasonication. Cell debris was removed from the lysates by centrifugation for 5 min at 1,500 x g for 5 min, and the soluble lysate was normalized to 30 µg protein lysate in 15 µL of Lysis Buffer using a Bradford protein concentration assay (Bio-Rad 5000006). Proteins were then reduced and alkylated by the addition of 10x tris(2-carboxyethyl)phosphine hydrochloride (TCEP; Sigma Aldrich C4706) and chloroacetamide (CAM; Sigma Aldrich C0267) prepared in 50 mM HEPES, pH 8.0, in MS-grade water for final concentrations of 10 mM and 40 mM, respectively. The samples were then diluted by the addition of CaCl_2_ (Sigma Aldrich C4901) in 50 mM HEPES, pH 8.0, for final concentrations of 0.9 M urea and 1 mM CaCl_2_. Digestion using 1:100 (w/w) protease to protein lysate with MS-grade Trypsin/Lys-C Protease Mix (Thermo A40009) was then performed by mixing overnight at 37 °C. The digested peptide samples were then acidified using a final concentration of 0.3% v/v MS-grade trifluoroacetic acid until pH < 3 by pH paper. The digested peptides were then desalted using SOLAµ™ SPE Plates (Thermo 60209-001), which were first activated with pure acetonitrile and then equilibrated with two washes of 0.1% TFA. The peptide samples were then loaded and washed three times with 0.1% TFA and one wash with 0.1% TFA before eluting twice with 70% acetonitrile and 0.1% FA in water. The desalted peptides were then completely dried in a centrifugal vacuum concentrator. The peptides were then reconstituted in MS-grade 0.1% FA in water.

### LC-MS/MS analysis

#### MOLT-4 proteomics

Data were collected using a TimsTOF Pro2 (Bruker Daltonics, Bremen, Germany) coupled to a nanoElute LC pump (Bruker Daltonics, Bremen, Germany) via a CaptiveSpray nano-electrospray source. Peptides were separated on a reversed-phase C_18_ column (25 cm x 75 µm ID, 1.6 µM, IonOpticks, Australia) containing an integrated captive spray emitter. Peptides were separated using a 50 min gradient of 2 - 30% buffer B (acetonitrile in 0.1% formic acid) with a flow rate of 250 nL/min and column temperature maintained at 50 °C. Data-dependent acquisition (DDA) was performed in parallel accumulation-serial fragmentation (PASEF) mode to determine effective ion mobility windows for downstream diaPASEF data collection^43^. The ddaPASEF parameters included: 100% duty cycle using accumulation and ramp times of 50 ms each, 1 TIMS-MS scan and 10 PASEF ramps per acquisition cycle. The TIMS-MS survey scan was acquired between 100 – 1700 *m/z* and 1/k0 of 0.7 - 1.3 V.s/cm^2^. Precursors with 1 – 5 charges were selected and those that reached an intensity threshold of 20,000 arbitrary units were actively excluded for 0.4 min. The quadrupole isolation width was set to 2 *m/z* for *m/z* <700 and 3 *m/z* for *m/z* >800, with the *m/z* between 700-800 *m/z* being interpolated linearly. The TIMS elution voltages were calibrated linearly with three points (Agilent ESI-L Tuning Mix Ions; 622, 922, 1,222 *m/z*) to determine the reduced ion mobility coefficients (1/K_0_). To perform diaPASEF, the precursor distribution in the DDA *m/z*-ion mobility plane was used to design an acquisition scheme for Data-independent acquisition (DIA) data collection which included two windows in each 50 ms diaPASEF scan. Data was acquired using sixteen of these 25 Da precursor double window scans (creating 32 windows) which covered the diagonal scan line for doubly and triply charged precursors, with singly charged precursors able to be excluded by their position in the m/z-ion mobility plane. These precursor isolation windows were defined between 400 - 1200 *m/z* and 1/k0 of 0.7 - 1.3 V.s/cm^2^.

#### Jurkat proteomics

The reconstituted desalted peptides were then analyzed using a nanoElute 2 UHPLC (Bruker Daltonics, Bremen, Germany) coupled to a timsTOF HT (Bruker Daltonics, Bremen, Germany) via a CaptiveSpray nano-electrospray source. The peptides were separated in the UHPLC using an Aurora Ultimate nanoflow UHPLC column with CSI fitting (25 cm x 75 µm ID, 1.7 µm C18; IonOptics AUR3-25075C18-CSI) over a gradient from 4% to 26% (v/v) of Mobile Phase B (MPB; 0.1% FA in ACN) in Mobile Phase A (MPA; 0.1% FA in water) over 60 min, then 26% to 32% MPB for 5 min, and finally 95% MPB for up to total 70 min with a flow rate of 350 nL/min with column temperature maintained at 50 °C. The TIMS elution voltages were calibrated linearly with three points (Agilent ESI-L Tuning Mix Ions; 622, 922, 1,222 m/z) to determine the reduced ion mobility coefficients (1/K_0_). diaPASEF was performed using the MS settings 100 m/z for Scan Begin and 1700 m/z for Scan End in positive mode, the TIMS settings 0.70 V⋅s/cm^2^ for 1/K_0_ start, 1.30 V⋅s/cm^2^ for 1/K_0_ end, ramp time of 120.0 ms, 100% duty cycle, ramp rate of 7.93 Hz, and the capillary voltage set to 1600 V. diaPASEF windows from mass range 226.8 Da to 1226.8 Da and mobility range 0.70 1/K_0_ to 1.30 1/K_0_ were designed to provide 25 Da windows covering doubly and triply charged peptides as confirmed by DDA-PASEF scans, whereas singly charged peptides were excluded from the acquisition due to their position in the m/z-ion mobility plane.

### Processing and statistical analysis of LC-MS/MS proteomics

The diaPASEF raw file processing and controlling peptide and protein level false discovery rates, assembling proteins from peptides, and protein quantification from peptides were performed using library free analysis in DIA-NN 1.8^44^. Library free mode performs an in silico digestion of a given protein sequence database alongside deep learning-based predictions to extract the DIA precursor data into a collection of MS2 spectra. The search results are then used to generate a spectral library which is then employed for the targeted analysis of the DIA data searched against a Swissprot human database (January 2021) for MOLT-4 proteomics or a Uniprot reviewed human database (April 2023).

#### MOLT-4 proteomics

Database search criteria largely followed the default settings for directDIA including: tryptic with two missed cleavages, carbamidomethylation of cysteine, and oxidation of methionine and precursor Q-value (FDR) cut-off of 0.01. Precursor quantification strategy was set to Robust LC (high accuracy) with RT-dependent cross run normalization. Proteins with low sum of abundance (<2,000 x no. of treatments) were excluded from further analysis and proteins with missing values were imputed by random selection from a Gaussian distribution either with a mean of the non-missing values for that treatment group or with a mean equal to the median of the background (in cases when all values for a treatment group are missing). Protein abundances were scaled using in-house scripts in the R framework (R Development Core Team, 2014) and resulting data was filtered to only include proteins that had a minimum of 3 counts in at least 4 replicates of each independent comparison of treatment sample to the DMSO control. Significant changes comparing the relative protein abundance of these treatment to DMSO control comparisons were assessed by moderated t test as implemented in the limma package within the R framework^45^.

#### Jurkat proteomics

Database search criteria largely followed the default settings for DIA-NN 1.8.1 ^44^ with the following modifications: “FASTA digest” and “Deep-learning-based spectra” settings enabled, “trypsin/P” (includes cleavage after P) for the protease with up to 1 missed cleavages carbamidomethylation of cysteine, oxidation of methionine, and N-terminal acetylation, and precursor Q-value (FDR) cut-off of 0.01, MBR enabled, and “Robust LC (high accuracy)” for precursor quantification strategy. The identified and quantified peptides from the DIA-NN analysis were further analyzed in R. Peptides were filtered by removing (i) reverse and contaminant peptides, (ii) peptides with Global.Q.Value (FDR for peptide across all samples) and/or PG.Q.Value (FDR for quantified protein for peptide) greater than 0.0105, and (iii) at least two observed intensities values for a given peptide in at least one condition. Protein intensities were then re-calculated using the MaxLFQ method provided in the DIA-NN R package(*66*) and then differential statistics was performed using the DEqMS (a modified LIMMA) R package(*67*) to determine p-value, fold change, and the Benjamini-Hochberg adjusted p-value.

### Reporter vectors

The Cilantro 2 reporter vector (PGK.BsmBICloneSite-10aaFlexibleLinker-eGFP.IRES.mCherry. cppt.EF1α.PuroR, Addgene 74450) was used for flow-based degradation assays. The following boundaries for Brd4 truncations were used: Brd4(BD1): (UniProt entry <u>O60885</u>, residues 14-138) or Brd4(BD2): (residues 306-418). Reporter cell lines were generated as described before^15^.

### BRD4 reporter stability analysis

K562-Cas9 cells expressing the Brd4(BD1)_eGFP_, Brd4(BD2)_eGFP_ or Brd4(BD1-BD2)_eGFP_ degradation reporters were resuspended at 0.7 × 10^6^ ml^−1^ and 50 µl of cell suspension was seeded in 384-well plates. Shortly after, cells were treated with DMSO (n=2) or drug (n=2) for 16 h. The indicated drugs were dispensed with a D300 digital dispenser (Tecan). The fluorescent signal was quantified by flow cytometry (FACSymphony flow cytometer, BD Biosciences). Using FlowJo (flow cytometry analysis software, BD Biosciences), the geometric mean of the eGFP and mCherry fluorescent signal for round and mCherry-positive cells was calculated. The ratio of eGFP to mCherry was normalized to the average of two DMSO-treated controls.

### Bison CRISPR screen for BRD4 stability

The Bison CRISPR library targets 713 E1, E2 and E3 ubiquitin ligases, deubiquitinases and control genes and contains 2,852 guide RNAs (Addgene #169942)^46^. Ten per cent (v/v) of the Bison CRISPR library was added to 10 × 10^6^ Brd4 (BD1)_eGFP_ or Brd4 (BD2)_eGFP_ K562-Cas9 cells and transduced (2,400 rpm, 2 h, 37 °C). Eight days later, cells were treated with drug or DMSO for 16 h and four populations were collected (top 5%, top 5–15%, lowest 5–15% and lowest 5%) on the basis of the Brd4_eGFP_ to mCherry mean fluorescent intensity (MFI) ratio on an MA900 Cell Sorter (Sony). Sorted cells were collected by centrifugation and subjected to direct lysis and amplified as described before^32^. Amplified sgRNAs were quantified using the Illumina NextSeq platform (Genomics Platform, Broad Institute). The screen was analysed by comparing stable populations (top 5% eGFP/mCherry expression) to unstable populations (lowest 5% eGFP/mCherry expression) as described before^32^.

### Chemical synthesis

Additional details are provided in the supporting information.

### Plasmids

The study utilized the following plasmids:

pcDNA5-Flag, Mint-Flag/HA (SFFV.BsmBICloneSite.Flag/HA.cppt.EF1α.PuroR), and Ivy-Flag/HA (SFFV.Flag/HA.BsmBICloneSite.cppt.EF1α.PuroR) for co-immunoprecipitation and DCAF16 mutant transduction. Mutant cell lines were generated as previously described^15^.

## Acknowledgements

This work was support by the National Institutes of Health (NIH) grants P01CA066996 and R35CA253125 (to B.L.E.), R01CA262188 and R01CA2144608 (to E.S.F.), R01CA218278 (to N.S.G. and E.S.F.), NIH High End Instrumentation grant (1S10OD028697-01) (to N.S.G.), the Howard Hughes Medical Institute (to B.L.E.), and departmental funds from Stanford Chemical and Systems Biology and Stanford Cancer Institute (to N.S.G.). Z.K. is supported by Swiss National Science Foundation (grant no. P500PB_214385). We thank Dr. Stephen M. Hinshaw and Dr. Roman C. Sarott for proofreading.

## Author contributions

Z.Z and W.S.B. contributed equally to this work. Z.Z., W.S.B., and N.S.G. conceived and initiated the study; Z.Z. designed and synthesized electrophilic degraders with the help of M.N.N. and J.Z.; W.S.B. designed and conducted cell biological experiments with the help of Z.J.; Z.K. and M.S. carried out Bison CRISPR screening; B.G.D., K.A.D., H.M.J., and D.M.A. performed whole-cell proteomics experiments; N.S.G., B.L.E., and E.S.F. supervised the project; The manuscript was written by Z.Z., W.S.B., and N.S.G. with input from all authors.

## Competing interests

N.S.G. is a founder, science advisory board member (SAB) and equity holder in Syros, C4, Allorion, Lighthorse, Voronoi, Inception, Matchpoint, CobroVentures, GSK, Shenandoah (board member), Larkspur (board member) and Soltego (board member). The Gray lab receives or has received research funding from Novartis, Takeda, Astellas, Taiho, Jansen, Kinogen, Arbella, Deerfield, Springworks, Interline and Sanofi. B.L.E. has received research funding from Novartis and Calico. He has received consulting fees from Abbvie. He is a member of the scientific advisory board and shareholder for Neomorph Inc., TenSixteen Bio, Skyhawk Therapeutics, and Exo Therapeutics. E.S.F. is a founder, scientific advisory board (SAB) member, and equity holder of Civetta Therapeutics, Proximity Therapeutics, and Neomorph, Inc. (also board of directors). He is an equity holder and SAB member for Avilar Therapeutics, Photys Therapeutics, and Ajax Therapeutics and an equity holder in Lighthorse Therapeutics. E.S.F. is a consultant to Novartis, EcoR1 capital, Odyssey and Deerfield. The Fischer lab receives or has received research funding from Deerfield, Novartis, Ajax, Interline, Bayer and Astellas. K.A.D. receives or has received consulting fees from Kronos Bio and Neomorph Inc. M.S. has received research funding from Calico Life Sciences LLC. All other authors declare no competing interests.

## Ethics declarations

## Additional information

